# Force generation of cardiomyocytes in engineered environments

**DOI:** 10.64898/2026.09.24.754056

**Authors:** Mangalika Sinha, Ulrike Rölleke, Ruben Haag, Malte Tiburcy, Johannes Blumberg, Ulrich S. Schwarz, Wolfram-Hubertus Zimmermann, Sarah Köster

**Affiliations:** Institute for X-Ray Physics, University of Göttingen, Germany; Cluster of Excellence “Multiscale Bioimaging: From Molecular Machines to Networks of Excitable Cells” (MBExC), University of Göttingen, Göttingen, Germany; Institute of Pharmacology and Toxicology, University Medical Center Göttingen, Göttingen, Germany; DZHK (German Center for Cardiovascular Research), partner site Lower Saxony, Göttingen, Germany; Institute for Theoretical Physics and Bioquant, Heidelberg University, Heidelberg, Germany

**Keywords:** hiPSC-derived cardiomyocytes, traction force microscopy, sarcomere imaging, force generation

## Abstract

Cardiomyocyte contraction is essential for the pumping action of the heart and deteriorates after myocardial damage, either as a consequence of irreversible cardiomyocyte injury or stiffening of the extracellular matrix, a process described as fibrosis. Cell geometry and substrate stiffness not only influence sarcomere architecture and contractility, they also determine how much work the cardiomyocytes can transfer to their environment. Human induced pluripotent stem cell (hiPSC)-derived cardiomyocytes provide a model system to examine these effects in a controlled manner, enabling us to mimic key features of the cardiac environment in vitro. Here we show that geometric confinement and substrate stiffness influence different aspects of cardiomyocyte force generation. By combining traction force microscopy and live-cell imaging of hiPSC-derived cardiomyocytes on soft and patterned substrates, we find that geometric confinement leads to longer resting sarcomeres and more pronounced sarcomere shortening on substrates with both physiological (10 kPa) and fibrotic (30 kPa) stiffness. Interestingly, the substrate stiffness itself has little effect on resting sarcomere length, but influences the peak contractile stress. On 10 kPa substrates, confined cells generate higher peak contractile stresses than unconfined cells, whereas this difference is not observed at 30 kPa. Confinement also results in faster mechanical relaxation at 10 kPa, with no detectable difference at 30 kPa. The combination of defined cell geometry and physiological stiffness leads to longer resting sarcomere lengths, more pronounced sarcomere shortening, higher peak contractile stress, and faster relaxation, features associated with a more mature cardiomyocyte phenotype. These findings show that the mechanical consequences of cell geometry depend on substrate stiffness and, therefore, both aspects have to be considered when employing hiPSC-derived cardiomyocytes as a model system in mechanobiology research, disease modeling and drug testing.

## Introduction

Cardiovascular diseases remain a major global health burden, highlighting the need for physiologically relevant human cardiac model systems to study disease mechanisms and to evaluate therapeutic strategies. Primary adult human cardiomyocytes remain a relevant reference model for cardiac function [1, 2], but their limited availability and the difficulties associated with their isolation and culturing restrict their use as experimental models [1]. Human induced pluripotent stem cell-derived cardiomyocytes (hiPSC-derived cardiomyocytes) offer an attractive alternative. However, hiPSC-derived cardiomyocytes typically exhibit an immature phenotype compared to adult cardiomyocytes, including differences in cellular morphology, sarcomere organization, metabolism, electrophysiology, and contractile function [3–6].

These limitations have motivated extensive efforts to promote hiPSC-derived cardiomyocyte maturation through biochemical, metabolic, electrical, geometrical and mechanical cues [7–9]. Among these, physical cues such as cell geometry and extracellular matrix (ECM) mechanics are important regulators of cardiomyocyte structure and function [10–14]. In hiPSC-derived cardiomyocytes, controlling cell shape and exposing cells to substrate stiffness within a physiological range have been shown to enhance contractile function and features associated with maturation [7, 15]. However, the relative contributions of cell geometry and substrate mechanics to different aspects of cardiomyocyte function remain incompletely understood.

Cardiomyocyte contraction is a multiscale process spanning molecular, sarcomeric, cellular and tissue levels. Sarcomere shortening generates forces that are transmitted through the cytoskeleton and cell-matrix adhesions to the surrounding ECM. Consequently, measurements of individual parameters, such as force generation [16] or sarcomere dynamics [17, 18] capture specific aspects of this complex process and provide complementary insights into the relationship between structural organization and mechanical output. Sarcomere organization is commonly been assessed using fixed-cell immunofluorescence [19–21]. Additionally, live-cell probes such as SiR-actin enable time-resolved visualization of the actin cytoskeleton and have been used to resolve myofibrillar structure and sarcomere dynamics [7, 22]. Fluorescent *α*-actinin reporter hiPSC-derived cardiomyocytes lines provide a sarcomere specific readout by labeling Z-discs, allowing for direct visualization and quantification of sarcomere dynamics in living cells [15, 17, 18]. Combined with traction force microscopy (TFM), such live cell imaging provides complementary views of cardiomyocyte structure and ability to generate forces [15]. However, because substrate stiffness, ECM composition, and geometric confinement vary widely across studies using these tools [7, 15], it remains difficult to isolate the individual contributions of these mechanical and geometric cues to cardiomyocyte structure and function.

Ribeiro et. al [7] showed that physiological cell shape and substrate stiffness together enhance the contractile function of patterned hiPSC-derived cardiomyocytes, while a later study [15] found that increasing substrate stiffness enhances contractile force but has little effect on sarcomere dynamics beyond the physiological range for cells on line patterns. For unconstrained embryonic cardiomyocytes on soft elastic substrates, it has been shown that contractile output peaks at the physiological stiffness of 10 kPa [10] and that cell shape, but not myofibril structure, depends on substrate stiffness [23]. Together, these findings suggest that the structural organization and force output may depend differently on the extracellular mechanical environment and cellular geometry.

Here, we study this hypothesis by systematically investigating how cell geometry and substrate stiffness jointly regulate the structural and mechanical function of hiPSC-derived cardiomyocytes. By comparing rectangular-shaped and unconfined cells on both physiologically relevant and fibrotic-like substrate stiffnesses, we examine how these cues influence cardiomyocyte structure and function. By applying a combination of TFM and live-cell sarcomere imaging to the same cells and under identical experimental conditions, we find that cell geometry and substrate stiffness influence different aspects of cardiomyocyte force generation. Sarcomere organization and dynamics are associated with cell geometry, whereas the substrate stiffness mostly influence the peak contractile stress and the cell-geometry-dependence of the relaxation kinetics. These findings highlight the importance of considering both cellular geometry and extracellular mechanics when developing cellular models of hiPSC-derived cardiomyocytes for applications such as disease modeling and drug testing.

## Experimental

### Fabrication of substrates

Glass-bottom dishes (#P35G-1.0-20-C, 35 mm diameter, No. 1.0 coverglass, 20 mm glass diameter; Mat-Tek, Ashland, MA, USA) are exposed to air plasma (0.4 mbar, 50 W; ZEPTO, Diener Electronics GmbH, Ebhausen, Germany) for 30 seconds. The plasma-activated surfaces are coated with 3-Aminopropyltrimethoxy silane (#281778, Sigma-Aldrich, St. Louis, Missouri, USA) and incubated at room temperature for 8 minutes. After silanization, the dishes are rinsed with ultrapure water and air-dried. Subsequently, they are incubated with 0.5% glutaraldehyde (#4157.1, Carl Roth GmbH & Co. KG, Karlsruhe, Germany; dissolved in Dulbecco’s phosphate buffered saline (DPBS); #D8537, DPBS; Merck) for 30 minutes at room temperature. After incubation, the dishes are again rinsed with ultrapure water and air-dried.

Fluorescent beads (#F8801, FluoSpheres® carboxylate-modified, diameter 0.1 *µ*m, red fluorescence (580/605); Thermo Fisher Scientific; Waltham, MA, USA) are passivated by mixing 2.4 *µ*L of the stock bead suspension with 157.6 *µ*L of ultrapure water and 20 *µ*L of PLL-g-PEG (1 mg/mL, PLL(20)-g[3.5]-PEG(2 kDa); SuSoS, Dübendorf, Switzerland). The mixture is sonicated three times for 5 minutes with 20 minutes break, and then centrifuged at 7.6 × g for 10 minutes. After a visible pellet forms, the supernatant is removed, and the bead pellet is resuspended in 100 *µ*L of 50 mM HEPES buffer (#391338, ≥99.5% titrated with NaOH to pH 8.0; Sigma-Aldrich). To prepare polyacrylamide (PAA) substrates with a final stiffness of 10 kPa or 30 kPa (as determined by rheology), acrylamide (#1610140, 40%; Bio-Rad Laboratories, Hercules, CA, USA), bis-acrylamide (#1610142, 2%; Bio-Rad Laboratories), and DPBS are mixed according to Table 1 and 5 *µ*L of passivated carboxylated beads are added.

**Table 1:** Concentrations of 40% acrylamide and 2 % bis-acrylamide solutions to prepare 10 mL PAA solution of different stiffnesses.

| Stiffness (kPa) | 2% Bis-acrylamide (mL) | 40% Acrylamide (mL) | DPBS (mL) |
| --- | --- | --- | --- |
| 10 | 1.5 | 1.25 | 7.25 |
| 30 | 0.75 | 2.5 | 6.75 |

Polymerization of PAA is initiated by adding 0.5 *µ*L ammonium persulfate (#1610700, 10% APS; Bio-Rad Laboratories) and 0.2 *µ*L TEMED (#1610801, Bio-Rad Laboratories) to the bead-PAA mixture. The final solution is mixed thoroughly and 15 *µ*L are pipetted onto circular glass coverslips (#41001115, diameter 15 mm; VWR Inc., West Chester, Pennsylvania, USA). The coverslips are immediately sandwiched with the pre-treated glass-bottom dishes, and the gels are polymerized upside down for 1 hour. After polymerization, the gels are soaked in DPBS and the top coverslips are gently removed. The gels are thoroughly rinsed with DPBS to remove unpolymerized monomers and stored submerged in DPBS until use for a maximum of two days. For unpatterned PAA gels, surface functionalization is achieved by coating the top layer with Sulfo-SANPAH (#A35395; 0.4 mM in 50 mM HEPES buffer, pH 8; Thermo Fisher Scientific). The gel is exposed to UV light (365 nm, two 8 W tubes; Herolab GmbH Laborgeräte, Wiesloch, Germany) for 8 minutes to activate the crosslinker. Following UV activation, 50 *µ*L of 0.1 mg/mL Synthemax solution (#3535; Corning, Arizona, USA) is applied to the gel surface and incubated overnight at 4°C to ensure uniform coating. The next day, the unpatterned gels are rinsed with DPBS and stored submerged in DPBS until use.

### Substrate patterning

We fabricate a master mold on a silicon wafer (2-inch, MicroChemicals, Ulm, Germany) using standard photolithography. Briefly, SU-8 3005 photoresist (micro resist technology GmbH, Berlin, Germany) is spin-coated to a height of 7 *µ*m, exposed to UV light through a lithography mask (Selba, Versoix, Switzerland) and developed. Polydimethylsiloxane (PDMS) elastomer (#5498840000, Sylgard 184, Biesterfeld Spezialchemie GmbH, Hamburg, Germany) is prepared at a 10:1 base–to–cross linker ratio. The PDMS mixture is degassed and poured onto the patterned silicon wafer placed in a Petri dish, degassed once more and cured at 65°C for 2 hours. After curing, the PDMS is carefully removed from the master wafer and cut into 1 cm2 square stamps. Before incubation in protein solution, the PDMS stamps are cleaned using ultrasound, first with isopropanol and then with ultrapure water. After drying, the stamps are exposed to air plasma (0.4 mbar, 50 W, ZEPTO) for 30 seconds. Subsequently, 100 *µ*L of 0.1 mg/mL Synthemax solution is applied to each stamp, and the stamps are incubated overnight at 4°C to ensure uniform coating. For validation of the pattern transfer process by direct visualization of the protein pattern, a mixture of 80% unlabeled fibronectin (#F1141, Merck, Darmstadt, Germany) and 20% rhodamine-labeled fibronectin (#FNR01, Cytoskeleton Inc., Denver, USA) (final protein concentration 0.1 mg/mL) is used instead of Synthemax.

On the following day, circular glass coverslips (diameter 15 mm; VWR) are cleaned using ultrasound using the same procedure as for the PDMS stamps. Once fully dried, the coverslips are plasma-treated for 4 minutes to roughen the surface so as to avoid bonding of the PDMS and the glass. Excess protein solution is carefully removed from the PDMS stamps using lint-free wipes. The moist stamps are gently placed face-down onto the plasma-treated coverslips. A 30 g weight is placed on top to ensure conformal contact. After 20 minutes, the weight is removed, and 100 *µ*L of DPBS is added on top of the patterned coverslips to prevent drying and preserve the pattern until transfer onto the PAA substrate.

As with the unpatterned substrates, polymerization of the PAA is initiated by adding 0.5 *µ*L of 10% ammonium persulfate and 0.2 *µ*L of TEMED to the bead–PAA mixture. The solution is mixed thoroughly, and 15 *µ*L are pipetted onto the patterned coverslips. The coverslips are immediately placed onto pretreated glass-bottom dishes, and the gels are allowed to polymerize upside down for 1 hour. A schematic of the complete substrate patterning process is shown in the supporting Material, Fig. S1.

### Cell culture

Cardiomyocyte differentiation of a hiPSC cell line (hiPSC-ACTN2-citrine-derived cardiomyocytes) [17] follows the protocol described by Tiburcy et al. [24]. We refer to these cells as hiPSC-derived cardiomyocytes. After thawing, hiPSC-derived cardiomyocytes are cultured in T25 cell culture flasks (#83.3910.002, Sarstedt, Nümbrecht, Germany) coated with Corning® Matrigel® growth factor reduced (GFR) Basement Membrane Matrix (#CLS354230-1EA, Merck). The Matrigel is diluted to a final concentration of 0.8% in ice-cold DPBS. 10 mL of the diluted Matrigel solution is added to the cell culture flask, and incubated at 37°C in a 5% CO_2_ atmosphere for at least 1 hour to allow complete polymerization, before use. Cells are maintained in serum-free Roswell Park Memorial Institute (RPMI) 1640 medium with GlutaMAX (#61870010, Gibco, Thermo Fisher Scientific), supplemented with 1% penicillin-streptomycin (#P06-07100, 10,000 U/mL penicillin and 10 mg/mL streptomycin in the stock solution; PAN-Biotech, Aidenbach, Germany) and 2% B27 supplement (#17504-044, Gibco, Thermo Fisher Scientific), hereafter referred to as “supplemented RPMI medium”. Cultures are kept at 37°C in a humidified atmosphere containing 5% CO_2_, and the medium is changed every second day for at least two weeks post-thaw.

For seeding onto PAA substrates, the cells are enzymatically dissociated using a digestion solution composed of StemPro® Accutase® (#A11105-01, Gibco, Thermo Fisher Scientific), supplemented with 0.025% Trypsin (#15090-046, Gibco, Thermo Fisher Scientific) and 20 *µ*g/mL DNase I (#260913, Merck). The digestion is performed for 20 minutes at 37°C. The reaction is stopped using supplemented RPMI medium containing 5 *µ*mol/L Y-27632 dihydrochloride (#orb154626, Biorbyt, Durham, NC, USA). The cells are seeded on micropatterned and unpatterned PAA substrates prepared in glass-bottom dishes. Each micropatterned gel (diameter 15 mm) is seeded with approximately 100,000 cells, while each unpatterned gel is seeded with approximately 5,000 cells. These values ensure a sufficient number of adhered cells that are well separated from each other on the gels. Cells are cultured for 18 hours in supplemented RPMI medium containing 5*µ*mol/L Y-27632. Subsequently, cardiomyocytes are maintained in supplemented RPMI medium without Y-27632, with medium changes every second day, at 37°C and 5% CO_2_ for 5 days before data acquisition.

### Imaging and traction force microscopy

We employ a sequential imaging approach to perform TFM combined with sarcomere dynamics analysis. The workflow begins with bright field imaging to document the cell morphology, followed by time-lapse acquisition of fluorescent bead displacement and, in a second step, fluorescently labeled *α*-actinin (referred to as sarcomere motion). Imaging is performed at a frame rate of 40 Hz for the patterned substrates and 30–40 Hz (in each case adjusted to the size of the imaged area) for the unpatterned substrates. All experiments are conducted on an inverted microscope (IX81, Olympus, Hamburg, Germany), equipped with a 60 × water immersion objective (UPlanApo, NA = 1.2, Olympus) and a CMOS camera (Orca Flash 4.0; Hamamatsu Photonics Deutschland GmbH, Herrsching am Ammersee, Germany). Cells plated on Synthemax-coated PAA substrates are placed in a stage-top incubation chamber (STX, Tokai Hit Co., Ltd., Fujinomiya, Shizuoka-ken, Japan), which maintains stable physiological conditions of 5% CO_2_ and 37°C throughout the experiment. All PAA substrates have a minimum thickness of 40 *µ*m to avoid an influence of the glass coverslip on the behavior of the cells, and we select regions with homogeneous bead coverage for imaging. We use a FITC filter set (excitation at 494 nm, emission at 537 nm; AHF Analysentechnik, Tübingen, Germany) to image *α*-actinin, and a Texas Red filter set (excitation at 572 nm, emission at 628 nm; AHF Analysentechnik) to image the embedded fluorescent beads.

### Data analysis

For the analysis of TFM data, we apply an optical flow method using the Kanade–Lucas–Tomasi (KLT) algorithm with a regularization-based approach [25]. First, the bead image sequences are downsampled from 16-bit to 8-bit in ImageJ [26]. We then use the Shi–Tomasi corner detector to identify up to 5000 bead positions, ensuring a minimum distance of 3 pixels between features to avoid double-counting of clusters. The pyramidal KLT algorithm tracks intensity gradients around the detected beads within a 64×64 pixel search window. Displacements are computed between successive images, and the full trajectory of each bead is reconstructed with respect to the first frame. We assume linear motion within each search window, using a forward Euler method for motion integration. To correct for global drift, we select a small, deformation-free region of the image sequence and subtract the detected drift from all calculated displacements. A Tukey window (*α* = 0.2) is applied to enforce a zero-velocity boundary condition at the edges of the displacement field.

Traction forces are calculated using fourier transform traction cytometry (FTTC) with Tikhonov regularization. The optimal regularization parameter is determined by a Generalized Cross-Validation (GCV) function [27–29]. The total force is given by

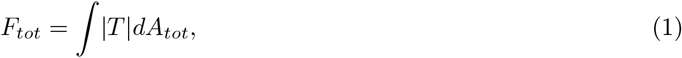

where *T* is the magnitude of traction vector at a given position and *dA*_*tot*_ defines the selected area containing a beating cardiomyocyte.

For analysis of the *α*-actinin image sequences, we first apply image deconvolution using the 2D-Richardson–Lucy algorithm with total variation regularization [30] and a calculated the Gibson-Lanni point spread function [31] of the microscope. The regularization parameter and the number of iterations are chosen manually to minimize the visible noise in the image. We use the *cupy* python package [32] to accelerate the calculation using a GPU. The deconvolved images are used to generate kymographs. From the kymographs, we determine the position of individual Z-discs as a function of time. These position traces are smoothed using a Savitzy-Golay filter (window length 51, polynomial order 3), and the time-dependent sarcomere length variation is calculated by subtracting the positions of adjacent Z-discs. The resulting sarcomere length traces are used for further analysis of sarcomere dynamics.

The relaxation phase of each contraction cycle is quantified by fitting the force decay to a single-exponential function

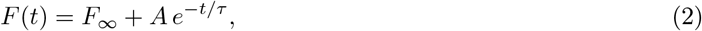

where *F* (*t*) is the traction force at time *t, F*_*∞*_ is the residual force after relaxation, *A* is the relaxation amplitude, and *τ* is the relaxation time constant. The fitting interval is selected automatically, with the fit starting at the point of maximum negative force derivative d*F/*d*t*. The model parameters are determined by non-linear least-squares fitting. All the analysis for sarcomere length changes over time and the relaxation time constant are performed using Jupyter notebooks (Python 3). The image analysis part for extracting the sarcomere length change from the *α*-actinin image sequences are carried out using the modules from Open Source Computer Vision (OpenCV) library [33]. The analysis of the frequency domain correlation of force and sarcomere are carried out using modules from Scientific Python (SciPY) library.

## Results

### In elongated cardiomyocytes traction forces are concentrated at the poles

To investigate the force generation of hiPSC-derived cardiomyocytes in different engineered environments, we perform TFM on patterned and unpatterned PAA substrates of two different stiffnesses, i.e., physiological 10 kPa and fibrotic 30 kPa. Fig. 1 provides an overview of the experimental procedure. Beating cardiomyocytes are cultured on Synthemax-coated PAA substrates with embedded fluorescent beads, enabling simultaneous visualization of substrate deformation by TFM and sarcomere dynamics through the fluorescently labeled Z-discs, see schematic in Fig. 1(a).

**Figure 1:**
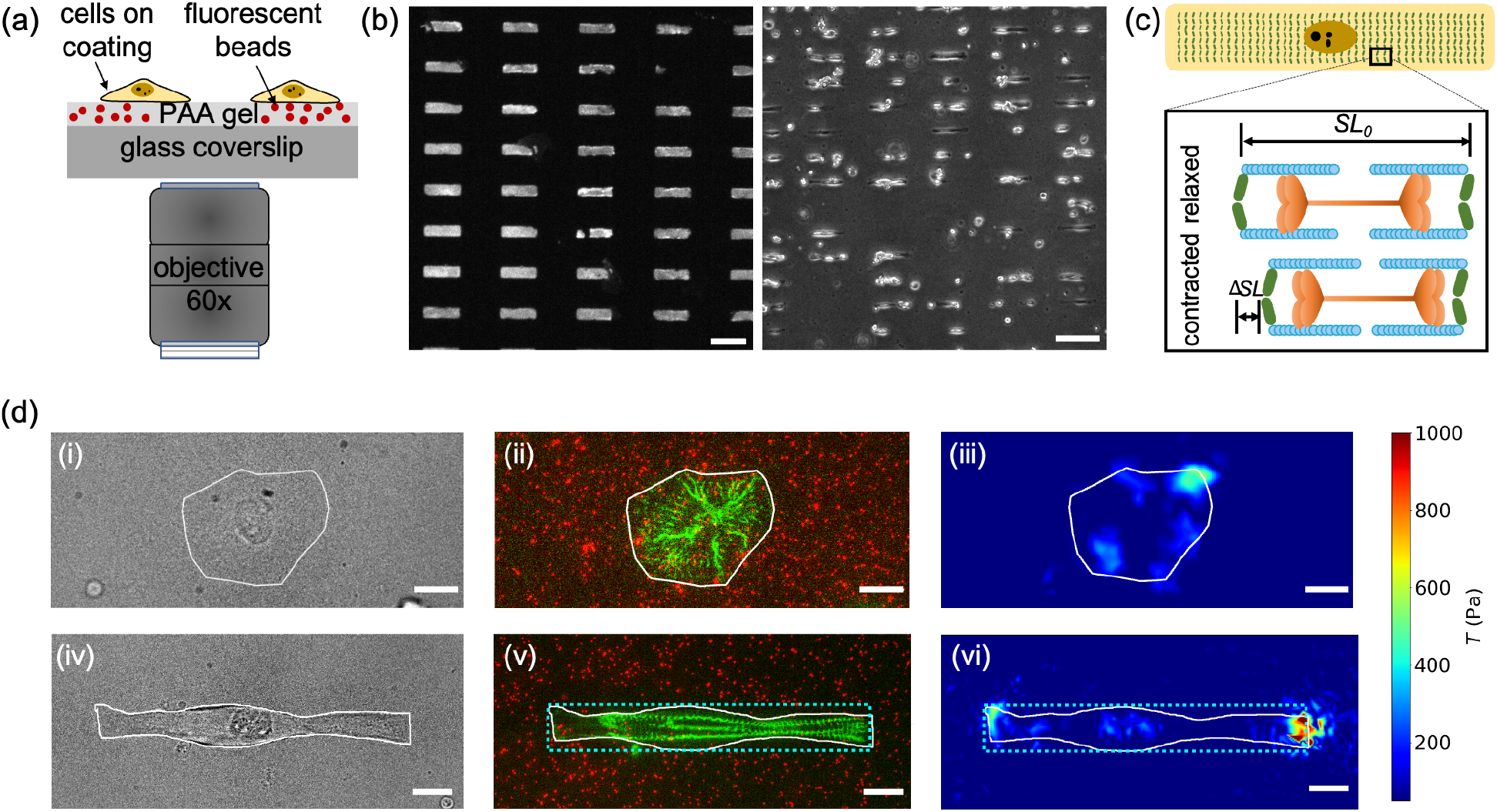
Traction force microscopy of cardiomyocytes. (a) Schematic of the experimental setup. Beating cardiomyocytes (yellow) are cultured on top of an elastic polyacrylamide (PAA) substrate (light gray, thickness 40–50 *µ*m) with embedded fluorescent beads (red). The substrate is coated with either rhodamine-labeled fibronectin (20%) for validation of the experiment or Synthemax for the measurements and imaged through a 60 × water immersion objective. (b) Patterned substrates and cell culture; (left) fluorescence image of rectangular fibronectin micropatterns on a PAA substrate; (right) phase-contrast image of cardiomyocytes adhering to micropatterns. Scale bars: 100 *µ*m. (c) Schematic of the sarcomere structure within a cardiomyocyte. Actin filaments (blue) and myosin filaments (orange) are anchored at Z-discs (green). The region between two adjacent Z-discs corresponds to a sarcomere. Upon contraction, the sarcomere shortens from its resting length *SL*_0_ by Δ*SL*. (d) Traction force microscopy of cardiomyocytes; (top) cardiomyocyte on an unpatterned substrate; (bottom) cardiomyocyte on a substrate with rectangular patterns (cyan boxes, 15 × 105 *µ*m); (left) bright field and (center) fluorescence images showing labeled Z-discs (green, *α*-actinin conjugated with citrine fluorescent protein) and the fluorescent beads (red); white outlines mark the cell boundaries; (right) corresponding traction stress maps, with the color scale indicating traction magnitude *T* . Scale bars: 10 *µ*m.

Fig. 1(b) shows a fluorescence image of patterned, fluorescently labeled fibronectin on a PAA substrate (left) and a phase-contrast image of cardiomyocytes cultured on a patterned substrate (right), confirming the high efficiency of the micropatterning strategy, as the majority of cells are confined to the adhesive rectangular patterns and adopt the intended geometry. Note that for validation of the pattern transfer process, we use fluorescently labeled fibronectin, whereas in all subsequent experiments, Synthemax is used as the coating material for both patterned and unpatterned substrates to provide a chemically defined and reproducible adhesive surface [34]. The patterning strategy is illustrated in the supporting material Fig. S1 and the details are provided in the Materials and methods section. The rectangular geometry of 15 *µ*m × 105 *µ*m is chosen to approximate the dimensions and aspect ratio (1:7) of adult human cardiomyocytes [6, 7].

Fig. 1(c) shows a schematic of the structural organization of the contractile Z-discs. The adjacent Z-dics define the boundaries of a sarcomere, i.e., the fundamental contractile unit of the myofibril. In the ACTN2-citrine cell line, *α*-actinin localized at the Z-discs is fluorescently labeled, allowing direct visualization of sarcomere organization and shortening during contraction. ATP-driven interactions between actin and myosin filaments shorten the sarcomere from its resting length, generating contractile forces that are transmitted to the substrate.

Fig. 1(d) shows representative TFM datasets of cardiomyocytes on an unpatterned (top) and a patterned (bottom) substrate. Bright field images reveal clear differences in morphology between the two culture conditions. On unpatterned substrates, cardiomyocytes spread into irregular and heterogeneous shapes, see Fig. 1(d,i), whereas on patterned substrates they consistently adopt elongated morphologies that follow the imposed geometry (Fig. 1(d,iv)). The corresponding overlays of the first frame of the fluorescent bead image and the consecutive ACTN2-citrine-labeled Z-dics image are shown in Fig. 1(d,ii,v), and the traction stress maps at the first peak of contraction reveal marked differences in force distribution, see Fig. 1(d,iii,vi) . While cardiomyocytes on unpatterned substrates generate a broad spatial distribution of traction stresses, cardiomyocytes on patterned substrates concentrate traction forces at the two cell poles, demonstrating that substrate patterning spatially organizes force generation along the major axis of the cell. The traction stress maps for both conditions are displayed using the same color scale, revealing higher peak traction stresses in patterned substrate, as evidenced by more pronounced red hotspots at the cell poles. The cell boundaries are indicated by white outlines in Fig. 1. For patterned cells, the imposed rectangular geometry is indicated by the cyan dashed outlines in Fig. 1(d,iv-vi). Additional examples of cardiomyocyte morphology on patterned and unpatterned substrates are provided in the supporting material Fig. S2. Time-lapse videos of the fluorescent bead displacement and *α*-actinin dynamics, together with the corresponding traction force maps for cardiomyocytes cultured on 10 kPa unpatterned and patterned substrates, are provided in the supporting material, movies S1 and S2.

### Substrate patterning enhances force generation on substrates of physiological stiffness

Having established that substrate patterning alters the spatial distribution of traction forces, we next investigate how cell geometry and substrate stiffness influence cardiomyocyte force generation. Fig. 2(a) shows typical force traces plotted against time for a single hiPSC-derived cardiomyocyte cultured on an unpatterned (magenta) and a patterned (cyan) substrate, illustrating the contractile behavior of the cells. Individual contraction peaks are identified automatically (black circles), and the peak force of each contraction cycle is extracted for further analysis (see supporting material Fig. S3(a)). To account for differences in cell spreading area, the peak force is normalized by the cell area (supporting material Fig. S3(b)), providing the peak stress *F*_peak_*/A*_cell_. Thus, we can directly compare the contractile stresses between cardiomyocytes of different geometries and sizes. The distributions of the peak stress are shown as violin plots in Fig. 2(b). On unpatterned substrates (magenta), cardiomyocytes cultured on the stiffer substrates exhibit significantly higher peak stress than those on physiological 10 kPa substrates, indicating that substrate stiffening enhances contractile force generation. This observation is consistent with previous studies on fibroblasts, including vinculin-deficient fibroblasts and other adherent cells, demonstrating that stiffer extracellular matrix environments promote increased cellular traction and contractility through enhanced force transmission at cell-matrix adhesions [35–39]. In contrast, on patterned substrates (cyan), the peak contractile stress decreases significantly from 10 kPa to 30 kPa, indicating that the effect of substrate stiffness on contractile stress depends on cell geometry.

**Figure 2:**
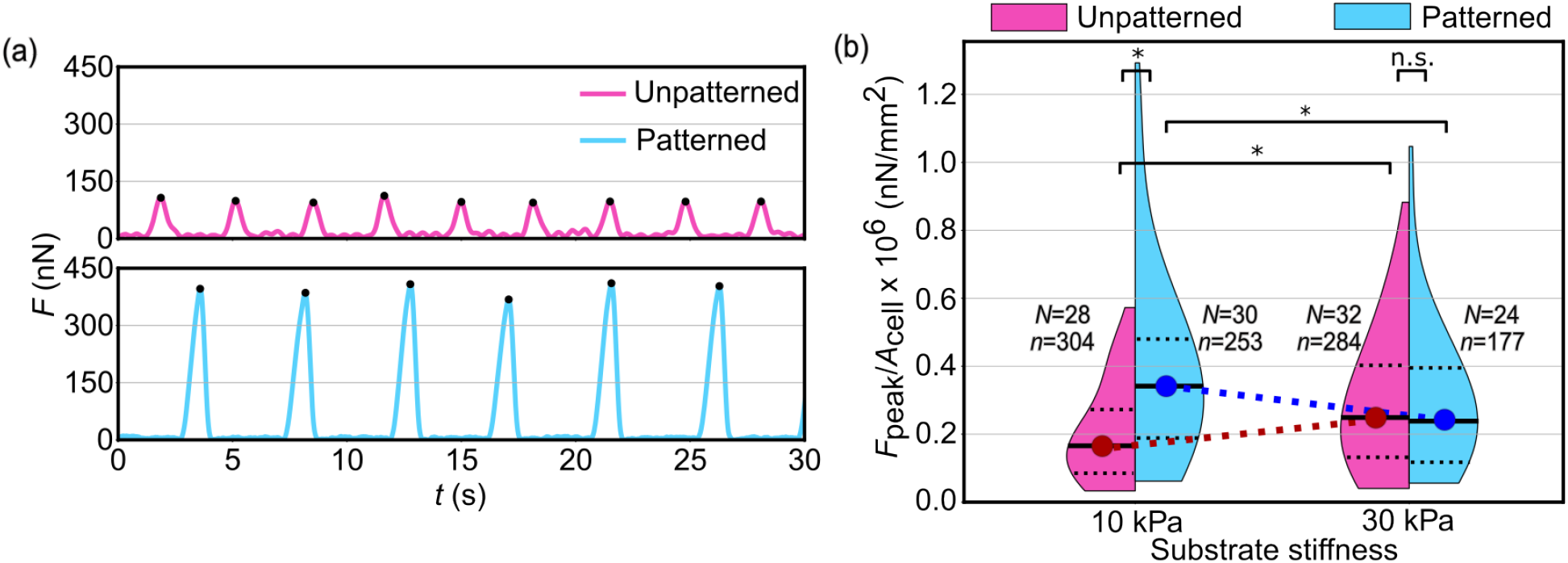
Contractile force generation of hiPSC-derived cardiomyocytes on different substrates. (a) Representative force traces of cells on unpatterned (magenta) and patterned (cyan) substrates over multiple beating cycles. The black dots mark the peak forces, *F*_peak_. (b) Distribution of the peak stresses *F*_peak_*/A*_cell_ for cells cultured on patterned and unpatterned substrates of different stiffnesses. The black solid lines represent the medians and the black dotted lines denote the interquartile range. The blue and red circles connected by dashed lines serves as visual guide to the eye, linking the median values of the patterned and unpatterned groups, respectively, across the two substrate stiffnesses. The sample sizes *n* indicate the number of analyzed contraction peaks, acquired from *N* cells per condition. Statistical comparisons are performed using two-sided Mann–Whitney U-tests (*: *p <* 0.05, n.s.: not significant).

When comparing elongated cells on patterned substrates (cyan) to randomly shaped cells on unpatterned substrates (magenta) for physiologically stiff (10 kPa) gels, we observe that the elongated cells generate significantly higher peak stresses. This finding demonstrates that geometric confinement promotes force generation under physiological stiffness. Interestingly, such a difference is not observed on 30 kPa substrates indicating that the beneficial effect of substrate patterning and induced cell elongation is not present on stiffer substrates. Together, these findings demonstrate that patterning enhances contractile force generation under physiological stiffness, whereas this advantage is lost on stiffer substrates, highlighting the role of substrate mechanics and cell geometry in regulating cardiomyocyte contractility.

### Substrate patterning enhances sarcomere shortening on substrates of physiological stiffness

Force generation in cardiomyocytes is a function of sarcomere alignment and contraction. We therefore next quantify the sarcomere organization of elongated and randomly shaped cells while varying the substrate stiffness. The analysis workflow is illustrated in Fig. 3(a). The raw fluorescence image sequence is computationally processed to reduce out-of-focus blur, and individual Z-discs are manually tracked (Fig. 3(a), left, see yellow line) to generate time-resolved Z-disc trajectories (Fig. 3(a), center). The temporal evolution of the sarcomere length (sarcomere traces) is calculated as the distance between adjacent Z-discs, *SL*_*t*_ = *Z*_*i*+1_(*t*) − *Z*_*i*_(*t*). Representative traces are shown for several neighboring sarcomeres in Fig. 3(a), right. For each sarcomere, the length trace is smoothed, and the minimum value within the upper 10% of the smoothed sarcomere-length distribution (dotted black lines, Fig. 3(a), right.) is defined as the resting sarcomere length *SL*_0_. This provides a robust estimation of the sarcomere length in the relaxed state.

**Figure 3:**
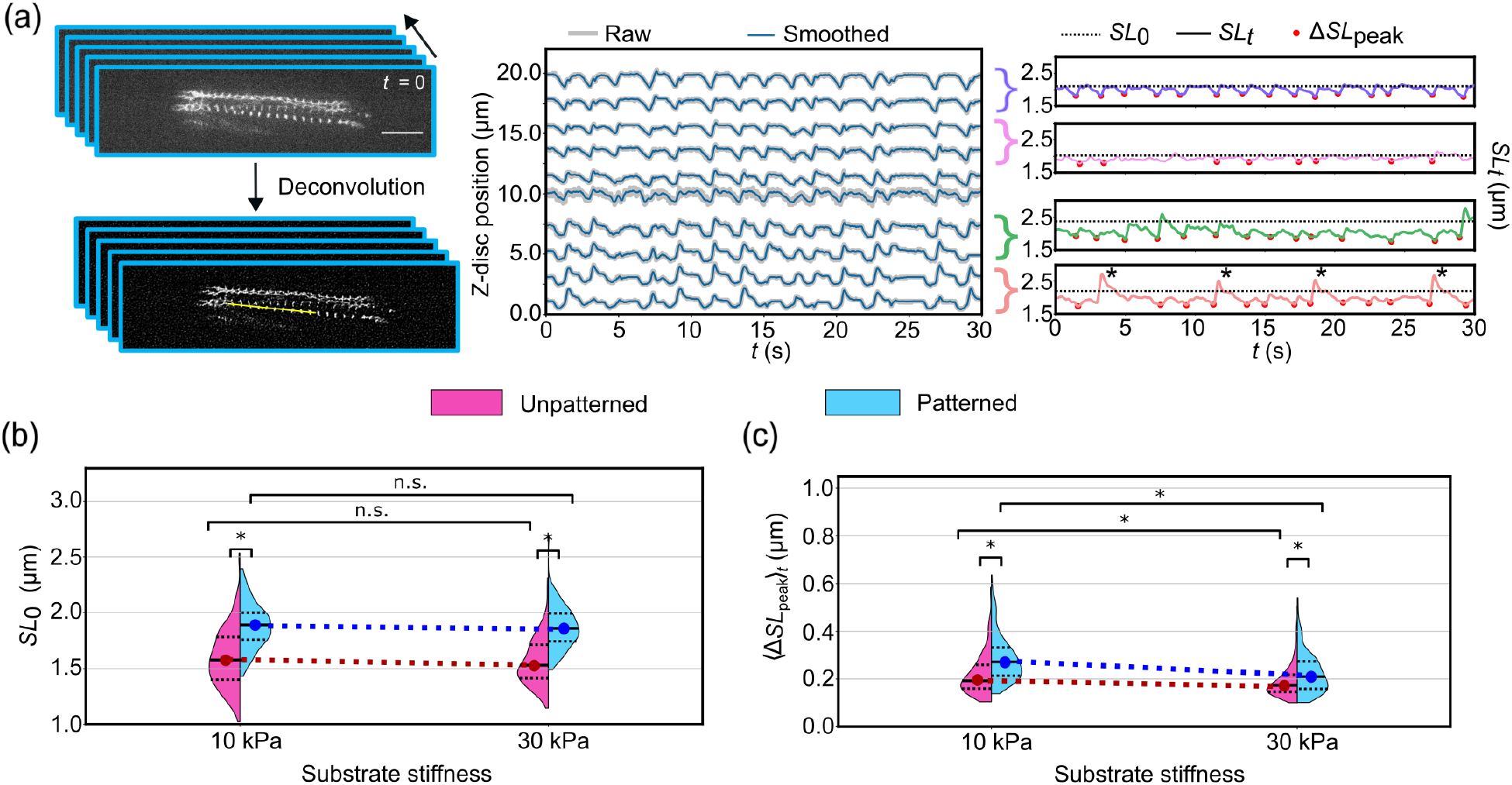
Quantification of sarcomere shortening dynamics of hiPSC-derived cardiomyocytes on different substrates. (a) Schematic of the analysis workflow for quantifying sarcomere shortening from fluorescence image sequences of individual hiPSC-derived cardiomyocytes. (Left) Raw fluorescence image stacks are first processed to enhance Z-disc structures, followed by manual identification of individual Z-discs (Scale bar: 10 *µ*m). (Center) Individual Z-discs are tracked over time to generate displacement traces. Gray lines represent the raw Z-disc traces over time and the blue lines show the corresponding smoothed traces. The distance between adjacent Z-disc pairs are then used to obtain sarcomere lengths as a function of time. (Right) Representative sarcomere length traces (*SL*_*t*_, 4 examples shown) obtained from the adjacent Z-disc pairs. For each *SL*_*t*_, the resting sarcomere length (*SL*_0_, dotted line) and local minima (red dots) are identified to determine the sarcomere shortening amplitudes, Δ*SL*_peak_. (b) Distribution of the resting sarcomere length *SL*_0_ for cardiomyocytes cultured on patterned (cyan) and unpatterned (magenta) substrates of different stiffnesses. (c) Distribution of individual sarcomere shortening amplitudes Δ*SL*_peak_ averaged over all contraction cycles recorded for a sarcomere Δ*SL*_peak *t*_. In both (b) and (c), the black solid lines represent the medians and the black dotted lines denote the interquartile range. The blue and red circles connected by dashed lines serve as visual guides to the eye, linking the median values of the patterned and unpatterned groups, respectively, across the two substrate stiffnesses. Statistical comparisons are performed using two-sided Mann–Whitney U-tests (*: *p <* 0.05, n.s.: not significant).

The sarcomere length change at each time point *t* is then calculated as Δ*SL*_*t*_ = *SL*_0_ − *SL*_*t*_. Contraction events are identified as local minima (dips) in the smoothed sarcomere-length traces, and the sarcomere shortening amplitude, Δ*SL*_peak_, is determined for each contraction cycle as the difference between *SL*_0_ and the sarcomere length at the detected minimum (red circles in Fig. 3(a), right). For each sarcomere, the resulting Δ*SL*_peak_ values are then averaged over all recorded contraction cycles yielding the time-averaged peak sarcomere shortening ⟨Δ*SL*_peak_⟩_*t*_. Occasional abrupt local increases in sarcomere length (see Fig. 3(a), right, black asterisks) are observed [17, 18]. As these events are not representative of the overall contractile behavior, they are excluded from the quantitative analysis.

The resting sarcomere length *SL*_0_ is a key structural parameter that reflects sarcomere organization. Immature hiPSC-derived cardiomyocytes typically exhibit resting sarcomere lengths of below 2 *µ*m [3–5], whereas maturation increases the resting sarcomere length towards 2.2 *µ*m, corresponding to the optimal overlap of actin and myosin filaments for maximal force generation [6]. The resting sarcomere lengths (*SL*_0_) as obtained from our analysis are shown in Fig. 3(b). Cardiomyocytes cultured on patterned substrates (cyan) exhibit consistently larger resting sarcomere lengths than cells on unpatterned substrates (magenta), indicating improved sarcomere organization in the elongated cell geometry. In contrast, within each patterning condition, the resting sarcomere length shows no significant dependence on substrate stiffness, suggesting that sarcomere organization is governed primarily by cell geometry rather than substrate stiffness.

Fig. 3(c) shows the distributions of the time-averaged peak sarcomere shortening values Δ*SL*_peak *t*_. On 10 kPa substrates, cardiomyocytes cultured on patterns (cyan) exhibit significantly larger Δ*SL*_peak *t*_ than cells on unpatterned substrates (magenta), demonstrating that the elongated cell geometry enhances contractile shortening under physiological stiffness. On 30 kPa substrates this is still the case, however, to a smaller extent, indicating that increased substrate stiffness diminishes the functional benefit provided by elongated cell geometry.

These findings demonstrate that substrate patterning regulates both the structural organization, reflected by the increased resting sarcomere length, and the contractile function of cardiomyocytes, reflected by the enhanced sarcomere length change amplitude. The substrate stiffness, by contrast, has little effect on resting sarcomere organization but predominantly modulates contractile function, reducing the advantage provided by elongated cell geometry under stiffer conditions.

### Relaxation times of elongated cells are shorter on substrates of physiological stiffness

We next quantify the relaxation time constant *τ* from the force traces obtained from TFM data by the workflow illustrated in Fig. 4a. Fig. 4a(i) shows that each contraction cycle consists of a rapid increase in force corresponding to the cardiomyocyte contraction, followed by a gradual decrease in force reflecting cardiomyocyte relaxation. *τ* is determined by fitting a single-exponential decay to the relaxation phase of each contraction cycle (see examples in 4a(ii,iii)). The exponential fit is started at the time point where d*F/*d*t* reaches its maximum, corresponding to the fastest rate of force relaxation [40]. *τ* is an important functional parameter that characterizes the rate at which cardiomyocytes return to their resting mechanical state following contraction. Shorter *τ* values correspond to more rapid mechanical relaxation and are generally associated with non-failing heart function [40, 41], associated with more efficient excitation-contraction coupling, faster calcium removal from the cytoplasm with simultaneous reuptake by the sarcoplasmic reticulum [42]. The *τ* values obtained from our analysis are shown in Fig. 4b. On 10 kPa substrates, cardiomyocytes cultured on patterned (cyan) substrates exhibit significantly shorter relaxation times than cells on unpatterned (magenta) substrates, indicating faster mechanical relaxation under physiological stiffness and confined cell geometry. In contrast, on 30 kPa substrates, relaxation times are comparable between cells cultured on patterned and unpatterned substrates. Since the spontaneous beating frequency can influence both contraction and relaxation kinetics [43], we additionally quantify the beating frequency under all four experimental conditions, see Fig. S4. The spontaneous beating frequency does not differ significantly between cells on patterned and unpatterned substrates for neither substrate stiffness. Thus, the shorter relaxation times observed for cells on patterned substrates at 10 kPa cannot be attributed to a difference in spontaneous beating frequency. We conclude that the benefit of substrate patterning on relaxation kinetics is not visible on stiffer substrates. These findings demonstrates that substrate patterning promotes faster mechanical relaxation only under physiological stiffness.

**Figure 4:**
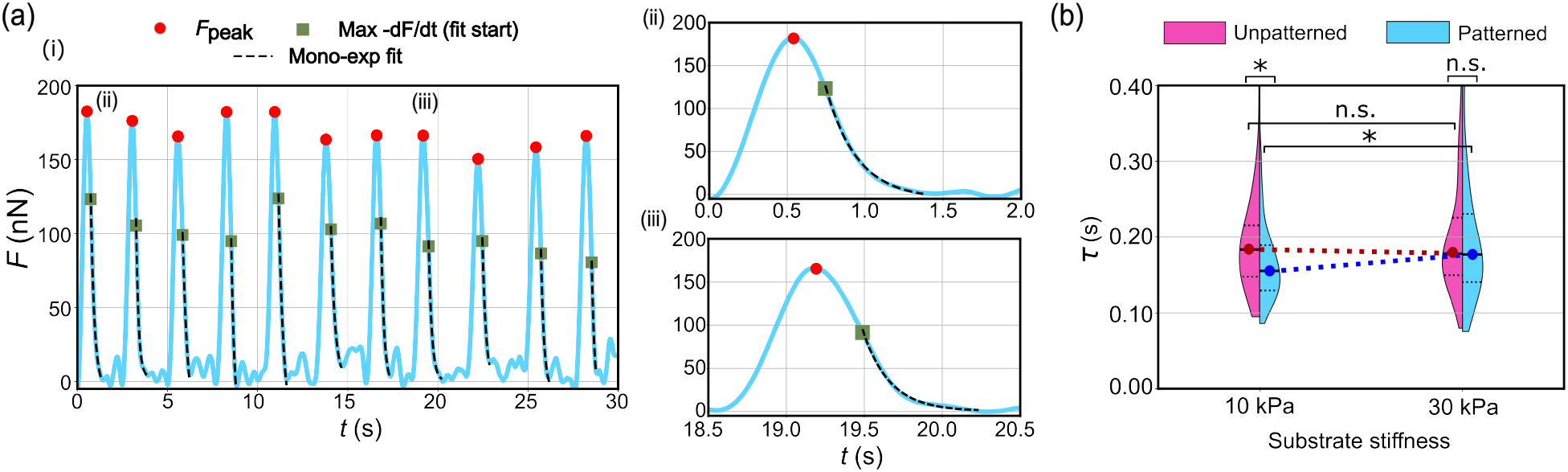
Relaxation kinetics of hiPSC-derived cardiomyocytes on different substrates. (a) Representative force traces illustrating the analysis workflow for relaxation kinetics. Force peaks (*F*_peak_) are identified for each contraction cycle (red circles) and the point of maximum relaxation rate (maximum −d*F/*d*t*, green squares) is used as the starting point for an exponential fitting (black dashed line) of the force decay to determine the relaxation time *τ* . Panel (i) and (ii) show representative exponential fits for individual contraction cycles. (b) Distribution of *τ* for cells cultured on patterned and unpatterned substrates of different stiffnesses. The black solid lines represent the medians and the black dotted lines denote the interquartile range. The blue and red circles connected by dashed lines serve as visual guide to the eye, linking the median values of the patterned and unpatterned groups, respectively, across the two substrate stiffnesses. Statistical comparisons are performed using two-sided Mann–Whitney U-tests (*: *p <* 0.05, n.s.: not significant).

## Discussion & Conclusion

Cardiomyocyte contraction involves coordinated changes in sarcomere organization, sarcomere dynamics, force generation, and finally relaxation that returns the cell towards its resting state and enables the next contraction cycle. In this study, we investigate how cell geometry and substrate stiffness influence these aspects of force generation in hiPSC-derived cardiomyocytes. We apply live-cell sarcomere imaging and TFM, sequentially to the same individual cells, thereby ensuring a detailed evaluation under identical experimental conditions. We find that geometric confinement is associated with longer sarcomere resting lengths at both investigated substrate stiffnesses, i.e. physiological (10 kPa) and fibrotic (30 kPa). This finding is consistent with previous studies demonstrating that cardiomyocyte shape and geometric constraint regulate sarcomere organization and myofibrillar alignment [7, 11, 12]. Geometric confinement also affects sarcomere dynamics, as cells on micropatterned substrates exhibit a more pronounced sarcomere shortening than unconfined cells. This is in line with previous observations that have shown controlled cell geometry to promote structural organization and functional maturation of hiPSC-derived cardiomyocytes [7, 11, 21, 44, 45]. Thus, geometric confinement is consistently associated with changes in both the resting organization and contractile behavior of the sarcomere.

In marked contrast to the effect of pattern geometry, substrate stiffness does not have a strong impact on sarcomere organization, in agreement with earlier reports for embryonic cardiomyocytes [23]. However, we observed that the effect of geometry on the peak contractile stress does vary with the substrate stiffness. Cells on physiologically relevant 10 kPa substrates with micropatterns exhibit higher peak contractile stress compared to unconfined cells. These higher peak stresses are not seen on 30 kPa substrates, in agreement with earlier reports for embryonic cardiomyocytes [10]. This difference suggests that a more pronounced sarcomere shortening does not necessarily correspond to a proportional increase in measured contractile stress. The stiffness-dependent changes in peak contractile stress suggest that the mechanical properties of the extracellular environment influence the force generated by the cell at the cell-substrate interface [10, 36– 39]. Together, these findings indicate that sarcomere shortening and peak contractile stress are not directly proportional and that the relationship between cell geometry and force output is shaped by the extracellular environment, specifically the substrate stiffness. However, the present experiments do not directly resolve the molecular or structural mechanisms underlying the stiffness-dependent differences in force output. Direct imaging and quantification of focal adhesions and other mechanotransduction components would be required to determine the molecular and structural mechanisms underlying the stiffness-dependent changes in force output [36–39].

In addition to contractile magnitude, geometry and substrate stiffness also influence the temporal behavior of contraction. Previous studies have shown that physiological cell geometry influences the contractile properties of patterned hiPSC-derived cardiomyocytes. Ribeiro et al. [7] compared rectangular micropatterns with aspect ratios of 1:1, 3:1, 5:1 and 7:1 geometry, and report the highest force generation in cells cultured on 7:1 rectangular micropatterns. In another study [15], they showed that increasing substrate stiffness enhances the contractility, whereas the sarcomere dynamics remains comparatively insensitive to stiffness above the physiological range in cardiomyocytes cultured on line patterns. The line patterns primarily guide cell alignment without strongly constraining the cell length, suggesting that the degree of geometric constraint may influence how extracellular environment affect cardiomyocyte contractility. Our findings extend these observations by showing that geometric confinement is also associated with changes in the temporal dynamics of contraction. Notably, the effect of geometry on relaxation is observed at 10 kPa but not at 30 kPa, indicating that the influence of cell geometry on relaxation kinetics is also dependent on extracellular environment, specifically substrate stiffness and is more pronounced at physiological stiffness.

In summary, we present an assessment of sarcomere organization, sarcomere dynamics, spatial force generation, and relaxation kinetics under controlled combinations of cell geometry and substrate stiffness. Interestingly, rather than affecting all aspects of cardiomyocyte function in the same manner, these mechanical cues appear to act at different levels of the contractile system: geometry strongly influences sarcomere organization and shortening, while substrate stiffness influences peak contractile stress and how cellular force output varies with cell geometry. The effects of cell geometry on the temporal behavior of contraction also depend on substrate stiffness, with faster relaxation observed for cells plated on substrates of physiological stiffness. Notably, combining defined cell geometry with physiologically relevant substrate stiffness (10 kPa) is associated with longer sarcomere resting lengths, greater sarcomere shortening, higher peak contractile stress, and faster relaxation in hiPSC-derived cardiomyocytes, features that have been associated with a more mature phenotype. Such control over the cellular environment provides a useful platform for studying cardiomyocyte mechanobiology and for developing in vitro models that more closely capture key features of the native myocardium, with potential applications in disease modeling and drug testing.

## Author Contributions

S.K. conceived and supervised the project. M.S. performed the experiments and the data analysis. U.S.S and J.B. developed the TFM data analysis procedure and code. R.H developed the deconvolution code used for the sarcomere analysis. M.T. and W.H.Z. provided the hiPSC cell line and associated protocols. U.R. performed the cell culture and assisted with sample preparation. M.S. and S.K. wrote the first draft of the manuscript, and all authors contributed to revising and finalizing the manuscript.

## Acknowledgements

We thank Daria Reher for technical support and helpful discussions. This work was financially supported by the German Federal Ministry of Research, Technology and Space (BMFTR) under grants No. 05K22MG3 and 05K25MGA (to S.K.) and the German Research Foundation (DFG): project-ID 449750155 - RTG 2756, projects A2, A7 (to S.K.) and under Germany’s Excellence Strategy - EXC 2067, ‘Multiscale Bioimaging: from Molecular Machines to Networks of Excitable Cells’ (MBExC, grant No. EXC 2067/1-390729940, to S.K. and W.H.Z.). W.H.Z. is additionally supported by the DZHK (German Center for Cardiovascular Research), the German Federal Ministry for Science and Education (BMBF FKZ 161L0250A), and the Foundation Leducq (20CVD04). J.B. and U.S.S. acknowledge support by the Carl Zeiss Foundation and the German Research Foundation (DFG) through the Cluster of Excellence “3D Matter Made to Order (3DMM2O)” (EXC-2082/1-390761711 and EXC-2082/2-390761711).

