## Supplementary Material for "Force generation of cardiomyocytes in engineered environments"

September 24, 2026

#### Supplementary figures

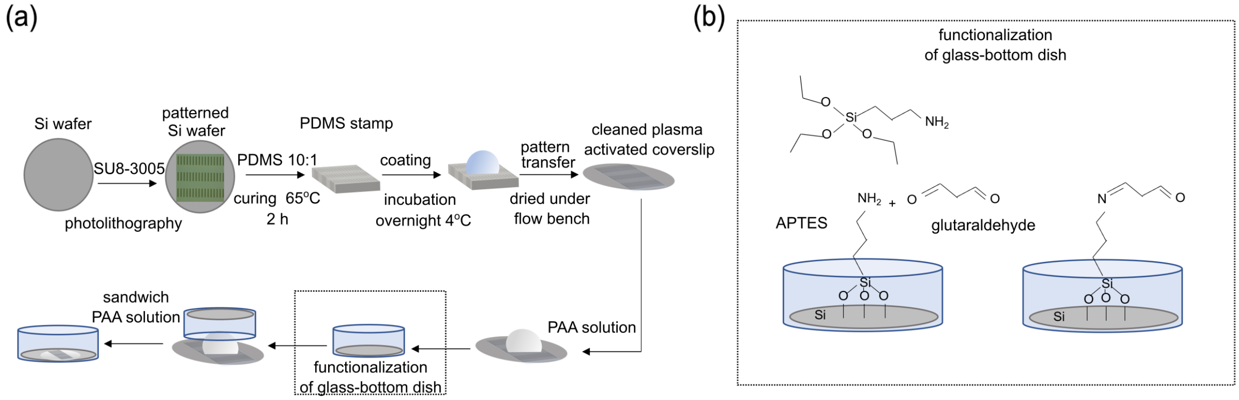


Fig. 1: Fabrication and functionalization of patterned substrates. (a) Schematic of the substrate fabrication workflow including photolithographic patterning of a silicon wafer, fabrication of a PDMS stamp, protein coating, pattern transfer onto a cleaned and plasma-activated glass coverslip, and subsequent polyacrylamide (PAA) polymerization. (b) Schematic of the glass-bottom dish functionalization, showing the surface modification with APTMS and glutaraldehyde to introduce reactive groups for PAA substrate attachment.

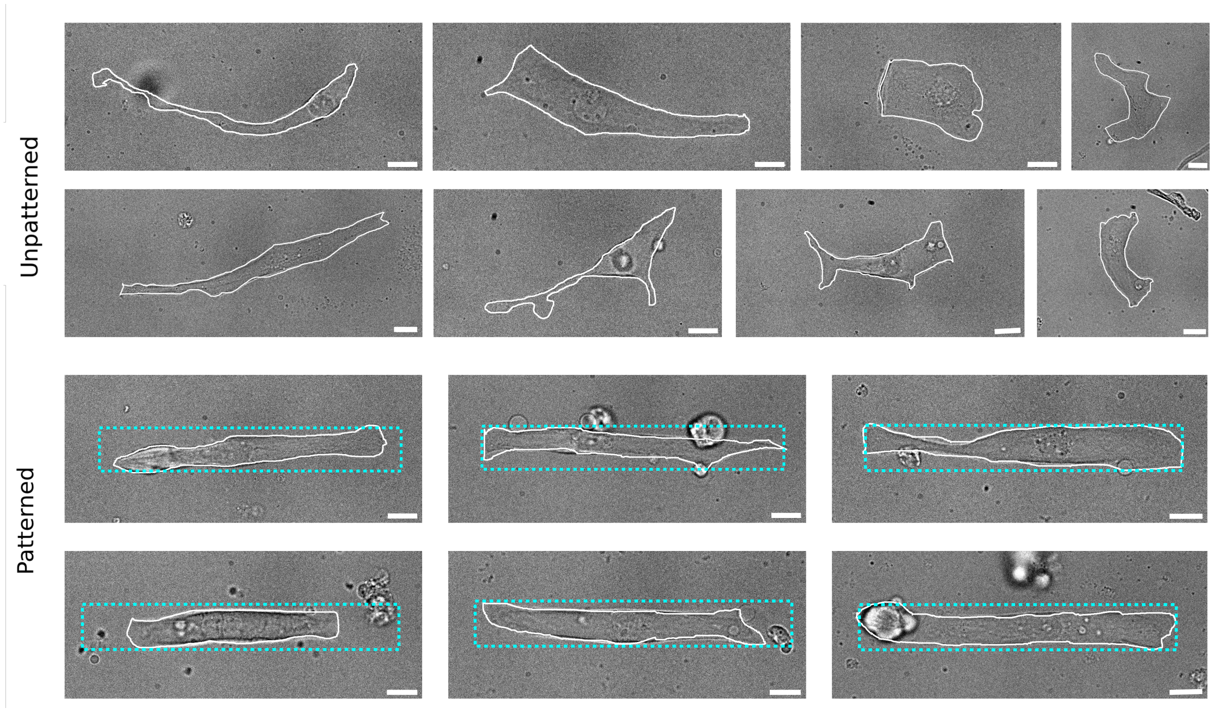


Fig. 2: Representative bright field images of cardiomyocytes cultured on unpatterned (top) and patterned (bottom)substrates. The white outlines marks the cell boundaries. The cyan boxes denote the dimension of the rectangular patterns (15 × 105 $\mu$m). Scale bars: 10 $\mu$m.

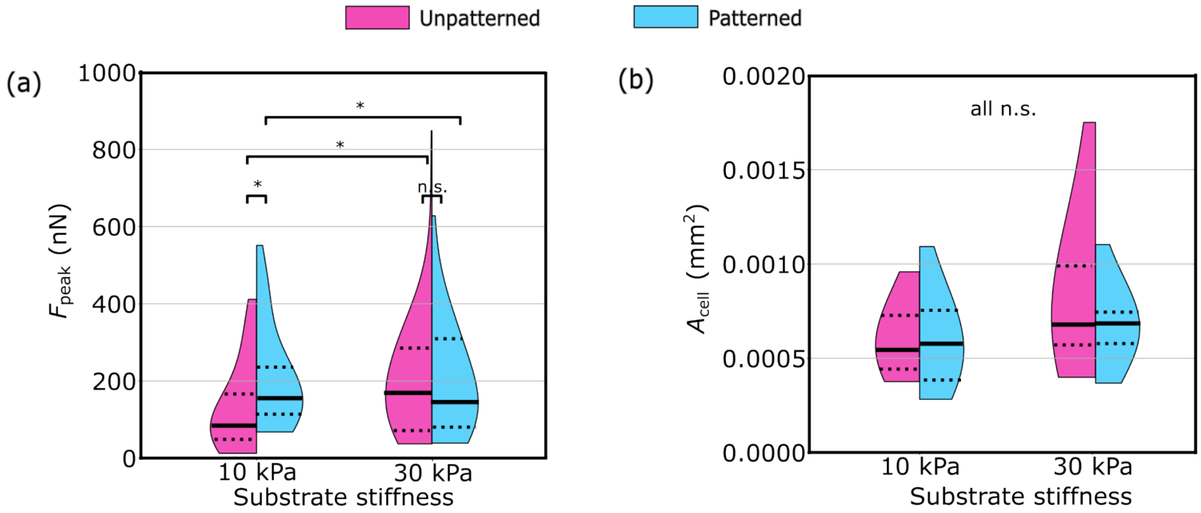


Fig. 3: Distributions of the (a) peak forces $F_{\text{peak}}$ and (b) cell areas $A_{\text{cell}}$ for the four investigated conditions. The black solid lines represent the medians and the dotted lines denote the interquartile range. Statistical comparisons are performed using two-sided Mann–Whitney U-tests. *: $p < 0.05$, ns: not significant.

### Supplementary movies

MOVIE S1: Time-lapse videos showing (left) fluorescent bead displacement, (center) the corresponding traction force, and (right) $\alpha$-actinin dynamics of an hiPSC-derived cardiomyocyte cultured on an unpatterned, 10 kPa substrate.

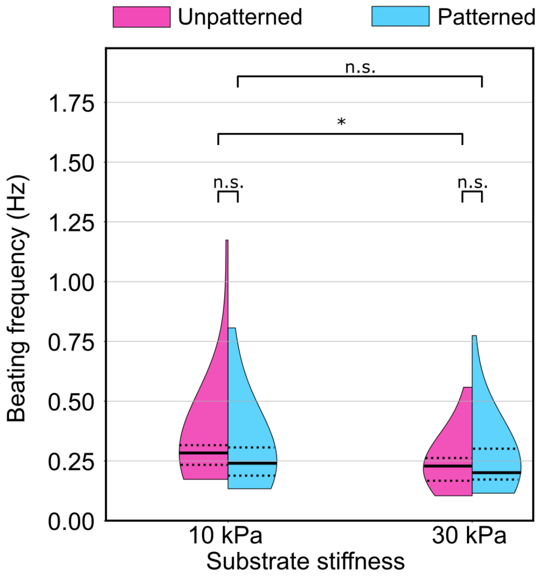


Fig. 4: Distributions of the spontaneous beating frequency for the four investigated conditions. The black solid lines represent the medians and the dotted lines denote the interquartile range. Statistical comparisons are performed using two-sided Mann–Whitney U-tests. *: $p < 0.05$, ns: not significant.

MOVIE S2: Time-lapse videos showing (left) fluorescent bead displacement, (center) the corresponding traction force, and (right) $\alpha$-actinin dynamics of an hiPSC-derived cardiomyocyte cultured a patterned, 10 kPa substrate.
